# CARD8 inflammasome sensitization through DPP9 inhibition enhances NNRTI-triggered killing of HIV-1-infected cells

**DOI:** 10.1101/2021.09.01.458624

**Authors:** Kolin Clark, Qiankun Wang, Liang Shan

## Abstract

Non-nucleoside reverse transcriptase inhibitors (NNRTIs) induce pyroptosis of HIV-1 infected CD4^+^ T cells through induction of intracellular viral protease activation, which then activates the CARD8 inflammasome. Due to high concentrations of NNRTIs being required for efficient CARD8 activation and elimination of HIV-1-infected cells, it is important to elucidate ways to sensitize the CARD8 inflammasome to NNRTI-induced activation. We show that this sensitization can be done through chemical inhibition of the CARD8 negative regulator DPP9. DPP9 inhibitor Val-boroPro (VbP) can act synergistically with NNRTIs to increase their efficacy in killing HIV-1-infected cells. We also show that VbP is able to partially overcome issues with NNRTI resistance and is capable of killing infected cells without the presence of NNRTIs. This offers a promising strategy for enhancing NNRTI efficacy in elimination of HIV-1 reservoirs in patients.

## Introduction

Despite the enhancement of combined antiretroviral therapy (cART) that allows people living with HIV-1 (PLWH) to have an undetectable viral load, there have only been two documented cases of complete remission from HIV-1 infection^1,2^. This clearly indicates the need for novel therapeutics for HIV-1 cure strategies. The primary hurdle in eradicating HIV-1 is the seeding of the latent reservoir which occurs quickly after infection^3^ primarily in activated CD4^+^ T cells that transition to resting memory cells^4^ and possibly in tissue macrophages^5^. These cells can self-replenish and evade all immune responses due to HIV-1 transcriptional inactivity^6^. However, in these latently infected cells, the integrated virus is still able to reactivate upon stimulation and spread infection^6^. This poses a significant barrier to HIV-1 eradication as current antiretroviral therapies prevent viral replication but do not remove the latent reservoir. One of the main strategies to eliminate the HIV-1 reservoir is through the “shock and kill” approach. This strategy utilizes latency reversal agents (LRAs) to reactivate the latent reservoir (shock) and then induce targeted cell death of infected cells (kill)^7^. Optimal efficiency is needed for both steps of this strategy, but we recently reported that the inflammasome sensor caspase recruitment domain 8 (CARD8) is able to sense intracellular HIV-1 protease activity and induce targeted cell killing of HIV-1 infected cells^8^.

The inflammasome is a multi-protein complex that is assembled upon sensing of their cognate ligand. Caspase-1 (CASP1) is the key effector for the inflammsome, and its active form can cleave Gasdermin D leading to pyroptosis. There are numerous pattern-recognition receptors (PRRs) that have been shown to activate the inflammasome and are characterized by either having a caspase recruitment domain (CARD) or a Pyrin domain (PYD) which can then in turn activate CASP1^9,10^. Recent studies demonstrated that one such PRR, CARD8, triggered the CASP1 activation and pyroptosis in human CD4^+^ T cells when cells were treated with the known Dipeptidyl Peptidase 9 (DPP9) inhibitor Val-boroPro (VbP) ^11, 12^. More recently, CARD8 was shown to sense intracellular HIV-1 protease activity^8^. CARD8 C-terminus (CARD8C) contains two key domains: the function-to-find domain (FIIND) and a CARD domain. Full-length CARD8 undergoes atuoprocessing at the FIIND domain leaving two non-covalently associated subunits^13^. HIV-1 protease was found to cleave CARD8 on the N-terminal subunit, which allows proteasomal degradation of the N-terminal fragment thereby freeing the C-terminal fragment^8^. The C-terminal fragment, in high enough concentrations, can then activate CASP1 and induce pyroptosis. However, freed C-terminal fragment may also be sequestered by the CARD8 negative regulator DPP9 which can inhibit pyroptosis efficiency^14^.

HIV-1 protease is not typically functional intracellularly before budding and it must be activated by other methods to be properly sensed by the CARD8 inflammasome. Premature intracellular protease activation can be achieved through the usage of non-nucleoside reverse transcriptase inhibitors (NNRTIs)^15^. This strategy offers benefits over other immune-based kill strategies that often rely upon recognition of the highly variable HIV-1 epitopes^16–18^ due to HIV-1 protease being less tolerant to mutation^19^. This is due to the critical need for the virus to maintain its enzymatic activities. Several reports have shown that NNRTIs, such as Efavirenz (EFV) and Rilpivirine (RPV), can induce HIV-1 protease-dependent killing of infected CD4^+^ T cells ^8, 20, 21^, which is due to activation of the CARD8 inflammasome^8^. It is clear that NNRTIs at micromolar concentrations drive Gag-Pol dimerization and intracellular protease activation which cleaves CARD8 leading to pyroptosis of HIV-1 infected cells. Strategies for enhancement of cell killing potency of these drugs are needed for efficient clearance of HIV-1-infected cells *in vivo*.

## Results

### NNRTIs induce death of HIV-1-infected cells in a dose-dependent manner

While NNRTI pharmacodynamics have been heavily studied for their ability to inhibit HIV-1 reverse transcription, they have yet to be studied in the context of their ability to activate the CARD8 inflammasome. To determine the *in vitro* pharmacodynamics of NNRTIs in CD4^+^ T cells, an HIV-1 reporter virus (pNL4-3-pol, see **Figure S1A**) was used to infect primary blood CD4^+^ T cells isolated from three independent healthy donors. Infected cells were then treated with Efavirenz (EFV), Rilpivirine (RPV), Etravirine (ETR), Doravirine (DOR), Nevirapine (NVP) in serial three-fold dilutions to assess the EC_50_ of killing for each NNRTI (**Figure 1A**). EFV, RPV, and ETR were able to induce robust cell killing at triple-digit nanomolar to low micromolar concentrations, whereas Doravirine and Nevirapine were ineffective at inducing cell death (**Figure 1B**). As macrophages are also key cellular targets for HIV-1, and were shown to have a functional CARD8 inflammasome^8, 22^, we also demonstrate a similar dose-dependent relationship (**Figure 1C)**. This relationship was due to CARD8 inflammasome activation as demonstrated by ablation of killing in *CARD8-KO* or *CASP1-KO* THP-1 cells. (**Figure 1D and E**).

**Figure 1:**
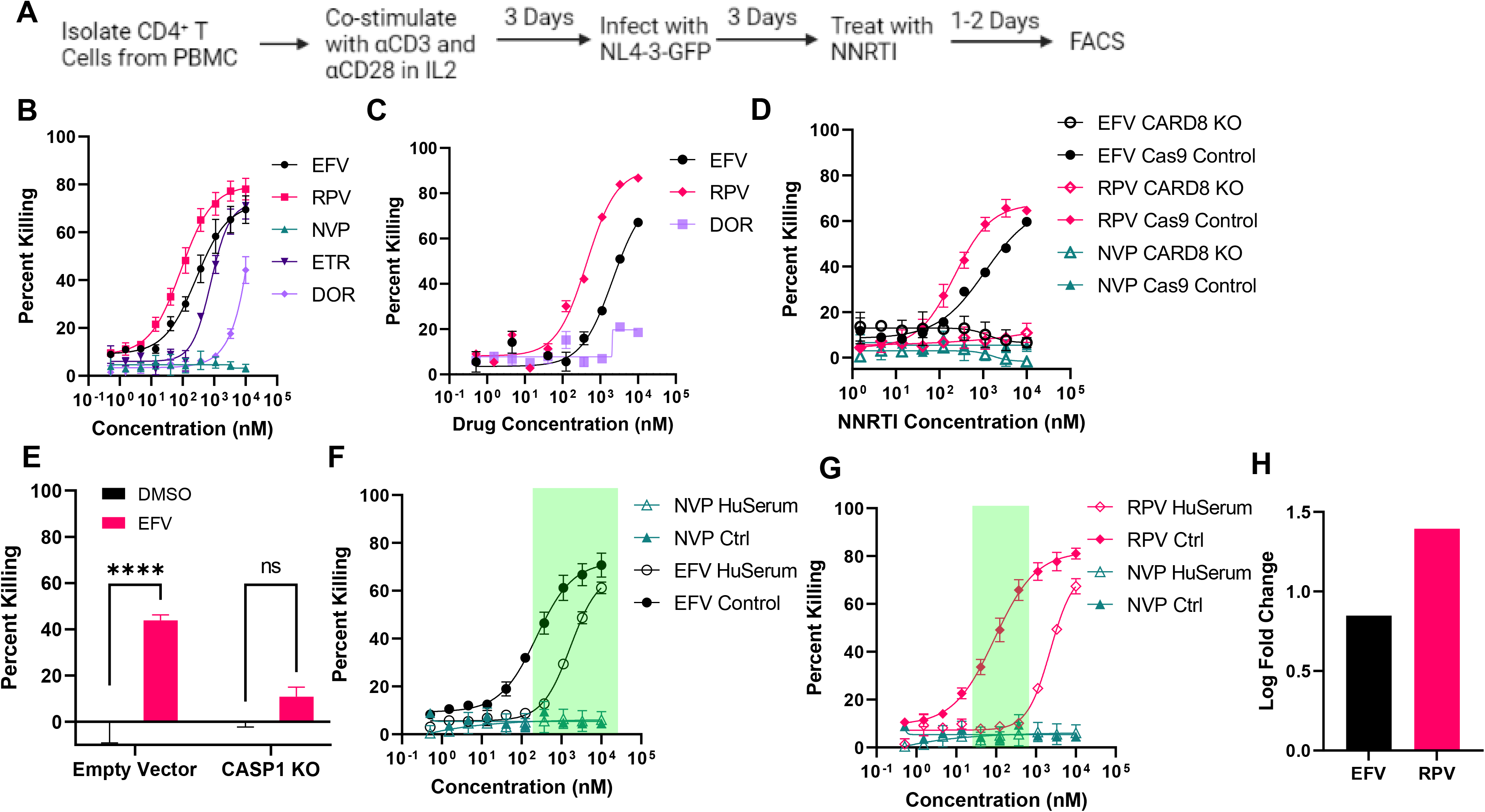
NNRTIs induce death of HIV-1-infected cells in a dose-dependent manner. **A)** Depiction of how CD4^+^ T Cells were treated with NNRTI’s and assayed for killing. **B)** Dose response curves for various NNRTI’s in successive three-fold dilutions in three healthy donor CD4^+^ T cells isolated from PBMC. EC_50_ Values for EFV, RPV, and ETR, are as follows: 266.1nM, 87.8nM, and 786.6nM respectively. ETR and DOR did not provide sufficient killing for EC_50_ calculation. **C)** Dose response curves for THP-1 infected with NL4-3-GFP and treated with EFV, RPV, or DOR as in panel A. EC_50_ for EFV is 2128nM and 438 for RPV. **D)** Dose response curves for killing of NL4-3-Δvif-vpr infected THP-1 *CARD8KO* or Cas9 transduced control cells treated with EFV, RPV, or NVP. The ability of EFV and RPV to kill infected cells is CARD8 dependent regardless of concentration. **E)** EFV based killing and activation of the CARD8 inflammasome is dependent upon CASP1 as evidenced by knockout of CASP1 in NL4-3-Δvif-vpr infected THP-1 cells with CRISPR/Cas9. (**** = p<.0001 by two-way ANOVA with Sidak’s multiple comparison test). **F and G)** Treatment of CD4^+^ T cells with EFV and RPV with or without the presence of 50% human serum in the culture media (NL4-3-pol). **G)** The log fold change increase in EC_50_ due to the presence of human serum.

The translatability of an NNRTI-based strategy for killing of HIV-1-infected cells is met with several barriers that can potentially reduce NNRTI efficacy *in vivo*. One key barrier to implementation is NNRTI’s high affinity for binding human serum proteins *in vivo^23^*. To assess this effect, CD4^+^ T cells were cultured with the presence of 50% human serum and show stark shifts in the dose response curves for both EFV and RPV (**Figure 1F and G).** With the presence of human serum, the dose response curve for RPV is shifted out of clinical concentration recommendations whereas EFV maintains some efficacy with the presence of human serum. Additionally, EFV is less affected by the presence of human serum as RPV as evidenced by a smaller log fold change in the EC_50_ (**Figure 1H**). These data suggest that EFV offers a distinct benefit over RPV for use in NNRTI based shock and kill strategies due to its higher plasma concentration and intracellular concentration^24, 25^. However, the efficacy of EFV is reduced with the presence of human serum which calls for the elucidation of strategies that could either increase intracellular NNRTI concentrations or sensitize the CARD8 inflammasome to NNRTI-based killing.

### DPP9 inhibition sensitizes the CARD8 Inflammasome to NNRTI-induced pyroptosis

DPP9 can bind to CARD8 first as a heterodimer with one copy of the full length CARD8 protein, then as a heterotrimer by catching a freed CARD8C^14^. As the C-terminal fragment is responsible for inflammasome activation, DPP9’s ability to catch CARD8C inhibits CARD8-induced pyroptosis. Overcoming DPP9 inhibition therefore should increase the rate of CARD8 inflammasome activation and sensitize the inflammasome to sensing HIV-1 protease activity. It was recently reported that VbP is able to bind to the DPP9-CARD8 heterodimer and prevent heterotrimer formation hence increasing intracellular CARD8C concentrations^14^. Additionally, VbP has another mechanism of action where it can induce N-terminal degradation of CARD8 which is also able to activate the inflammasome, although the direct mechanism of action has yet to be elucidated^14, 26^. We therefore posited that VbP’s ability to inhibit DPP9 may act synergistically with NNRTI-based killing due to sensitization of the CARD8 inflammasome (**Figure 2A**).

**Figure 2:**
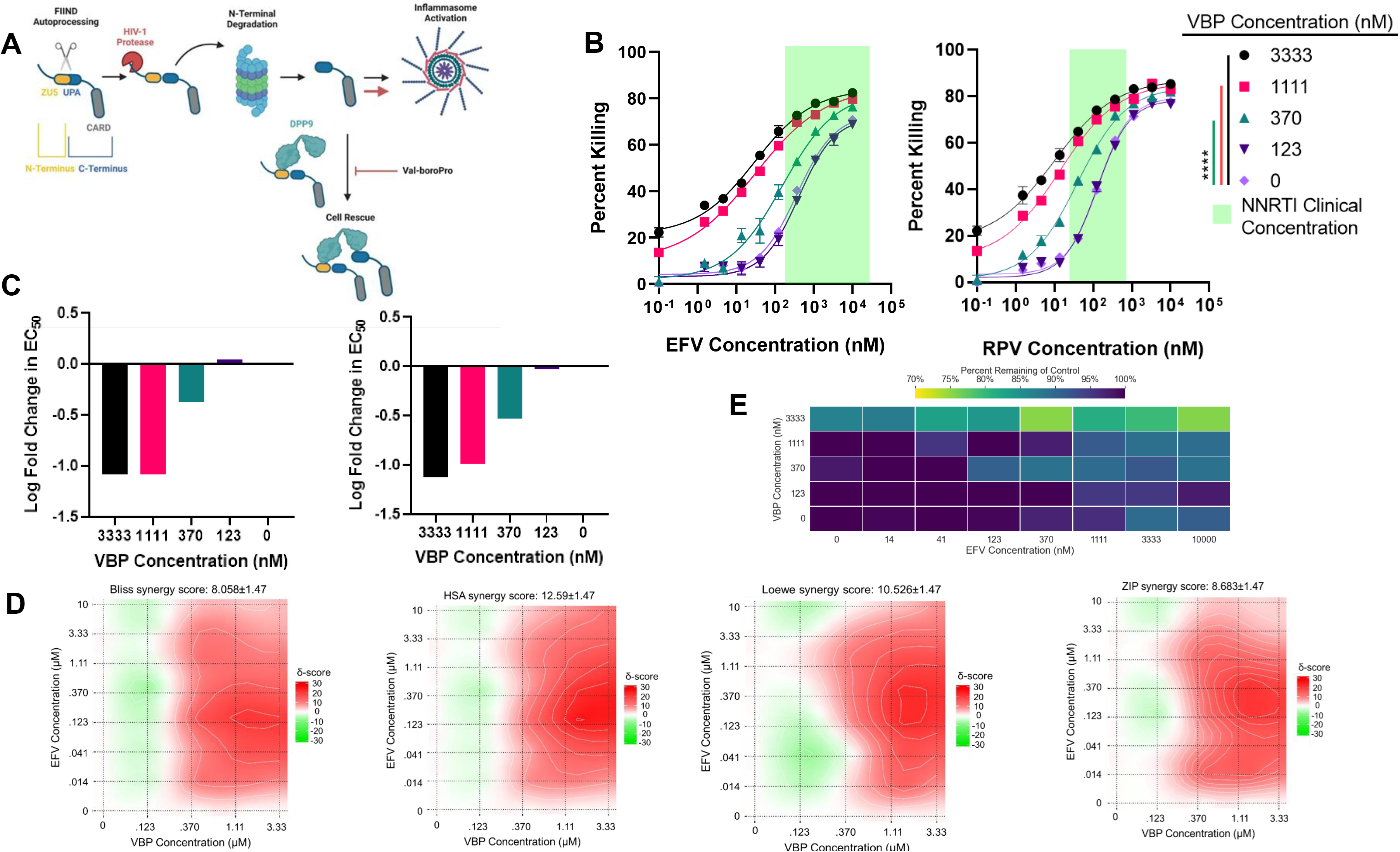
DPP9 inhibition sensitizes the CARD8 Inflammasome to NNRTI-induced pyroptosis. **A)** Graphical depiction of CARD8 inflammasome activation and sensitization through VBP. CARD8 first undergoes autoprocessing of its FIIND domain leaving two non-covalenetly bound fragments (N-terminal and C-terminal). HIV-1 protease, after premature activation due to Gag-Pol dimerization through NNRTIs, can cleave the N-terminal fragment leading to proteasomal degradation. The released C-terminal fragment can either activate the inflammasome through CASP1 or be caught by DPP9. VBP inhibits DPP9 and allows more freed C-terminal fragment to activate the inflammasome. **B)** Dose response curves for three donors of primary CD4^+^ T cells treated with EFV or RPV in combination with VBP. The green highlighted area denotes the drug plasma concentration range. Zero NNRTI concentration values were plotted at 10-1 nM concentration to allow log-transformation (****= p<.0001 by two-way ANOVA with Tukey’s multiple comparison test). **C)** Log fold changes in EC50 due to varying VBP concentrations are plotted for EFV (left) and RPV (right). **D)** VBP and EFV combination treatment denotes a synergistic relationship. Four independent synergy calculation methods from SynergyFinder2.0 (Bliss, HSA, Loewe, and ZIP) were used. **E)** VBP toxicity in CD4^+^ T Cells as denoted by the heatmap of MTS assay results of three separate donors of primary CD4^+^ T Cells treated for two days with EFV and/or VBP.

As expected, NNRTI induced killing of HIV-1 infected CD4^+^ T cells was enhanced upon treatment with VbP (**Figure 2B**). This enhancement of NNRTIs was shown to be dose-dependent upon increasing concentrations of VbP. Upon addition of VbP, the EC_50_ had log fold change shifts up to −1.1 for both EFV and RPV (**Figure 2C)**. This has the potential to overcome the EC_50_ shift due to the presence of human serum and demonstrates that DPP9 inhibition is essential for CARD8 inflammasome activation *in vivo*. Due to VbP’s ability to inhibit the capture of CARD8C by DPP9, we hypothesized that this relationship would be synergistic in nature. To further understand this complex relationship, we used SynergyFinder2.0 to identify whether this relationship was additive or synergistic^27^. As expected, combination treatment of VbP with EFV or RPV was found to be synergistic by four synergistic modeling methods: HSA^28^, BLISS^29^, Loewe^30^, and ZIP^31^ (**Figure 2D and S2A**). To evaluate the non-specific killing, we tested VbP in both HIV-1-infected and uninfected cells. VbP at concentrations lower than 3.33 μM has no significant toxicity in uninfected CD4^+^ T cells, indicating this mechanism of cell killing is specific to HIV-1 (**Figure 2E, S2B, and S2C)**. Additionally, VbP is able to induce low levels of cell killing of HIV-1 infected cells (22% and 13% for 3.33 μM and 1.11 μM respectively) (**Figure 2B**). We hypothesize that this may be due to low levels of spontaneous intracellular dimerization of gag-pol in infected cells which is insufficient to induce killing but becomes sufficient upon sensitization of the CARD8 inflammasome by VbP.

### Characterization of VbP enhancement of NNRTI-induced cell killing

To first begin understanding the dynamics of VbP enhancement of NNRTI induced cell killing we analyzed killing in CD4^+^ T cells upon combination or single treatment across time. Upon treatment with EFV at physiologically relevant concentration, the killing of HIV-1-infected cells became more rapid and robust with the presence of VbP (**Figure 3A**). It was previously shown that inhibition of the CARD8C capture by DPP9 was a rapid response, which we hypothesize is the main contributor to rapid enhancement of NNRTI induced cell killing^14^. There is a second phase of cell killing between 6 and 24 hours before the maximal killing plateaus for both EFV and combination treatments. When looking at the cellular killing by combination treatment in comparison to EFV alone, the fold change enhancement remains relatively consistent indicating a rapid but uniform enhancement across time (**Figure 3B**). In VbP only group, killing was not found until 48 hours post treatment. This slow and low level killing could either be due to the levels of CARD8C generated by inefficient spontaneous Gag-Pol dimerization now being sufficient with DPP9 inhibition or N-terminal degradation of CARD8 directly induced by VbP adding to the pool of HIV-1 PR-cleaved CARD8 fragments thereby inducing the inflammasome activation. This experiment was repeated for THP-1 cells and demonstrated similar results to CD4^+^ T Cells (**Figure 3C and D**).

**Figure 3:**
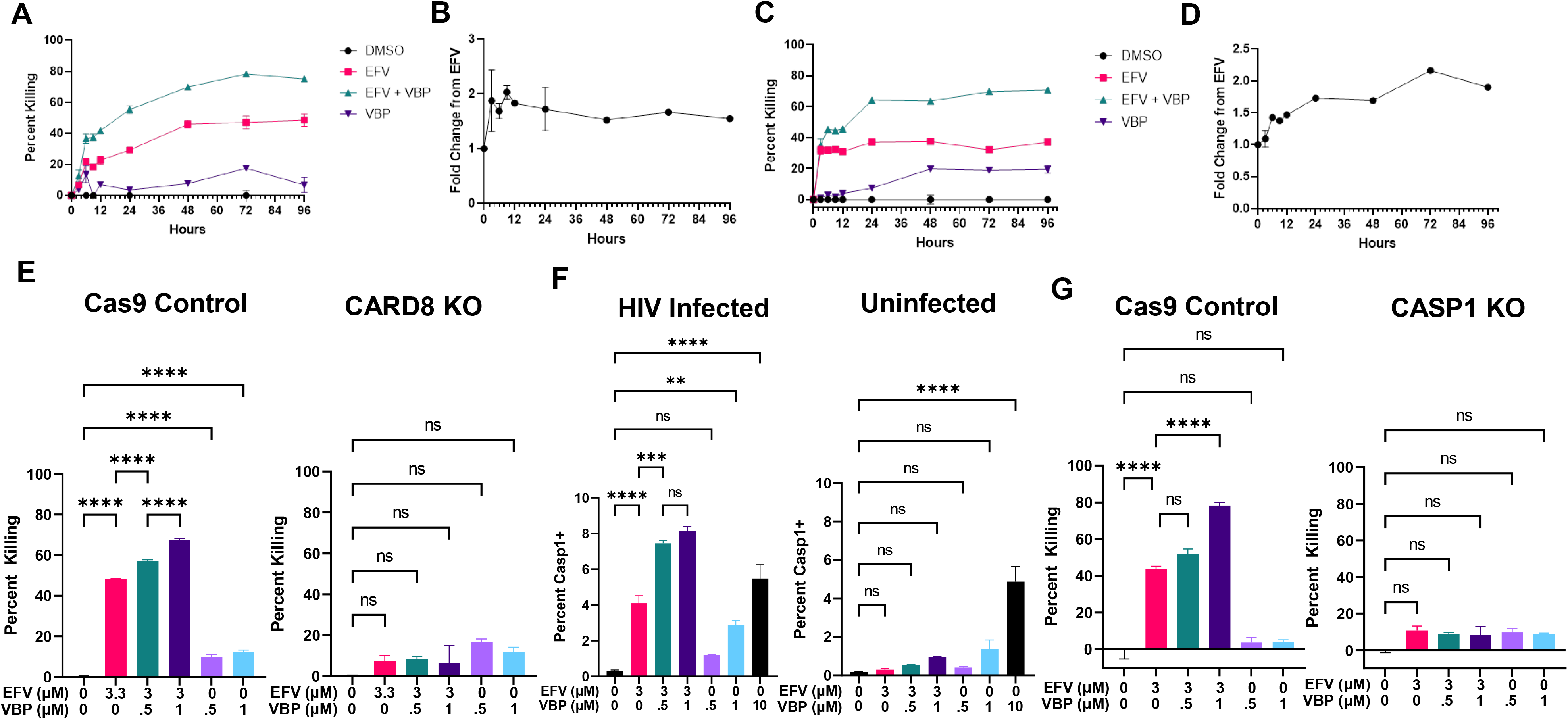
Characterization of VBP enhancement of NNRTI-induced cell killing. **A)** Time course treatment of HIV-1-infected primary CD4^+^ T cells treated with DMSO, EFV (.5μM), VBP (.5μM), or combination. Killing of infected cells plateaus after 48 hours in all conditions. **B)** Fold change enhancement of combination treatment in comparison to EFV alone treatment from panel A. **C and D)** Time course treatment and fold change enhancement of HIV-1-infected THP-1 cells treated with DMSO, EFV (1μM), VBP (.5μM), or combination. **E)** NNRTI-based killing and VBP enhancement is specific to the CARD8 inflammasome. **F)** CASP1 activation by NNRTIs and VBP. Cells were simultaneously treated with EFV and VBP and stained with CASP1 staining dye. **G)** CASP1 is required for the cell killing by NNRTIs and VBP (** = p<.01, *** = p<.001, **** = p<.0001 by one-way ANOVA with Tukey’s multiple comparison test).

To eliminate the possibility that VbP-based enhancement of NNRTIs is due to an unknown mechanism of cell death, we used *CARD8*-KO THP-1 cells and tested combination treatment in comparison to Cas9 control cells. Upon combination treatment in *CARD8*-KO cells, all killing was abolished when treated with EFV alone and the combination (**Figure 3E**). This clearly shows that any additional killing by the incorporation of VbP to NNRTI treatment is dependent upon CARD8 for its mechanism of action. As NLRP1 is also known to bind to DPP9 which can be released by VbP, we generated *NLRP1*-KO THP-1 cells and showed that enhancement was still present, which exclude the possibility that VbP enhancement is dependent on or regulated by NLRP1 (**Figure S3A**)^32,33^. Additionally, VbP is known to bind to both DPP9 and DPP8, we therefore show that knock-down of DPP8 does not ablate NNRTI enhancement suggesting a DPP9 specific mechanism of action (**Figure S3B**) ^34^. To ensure that the downstream components of the CARD8 inflammasome were demonstrating enhanced activation, we infected primary CD4^+^ T cells and treated the cells concurrently with a dye that specifically stains the active form of CASP1. As can be seen in **Figure 3F**, addition of VbP to EFV showed increased CASP1 activation in HIV-1-infected CD4^+^ T cells which is not the case for uninfected cells. Additionally, VbP alone at low concentrations (≤1 μM) shows significant CASP1 activation specifically in HIV-1-infected cells, suggesting that it relies upon the presence of HIV-1 to help induce the CARD8 inflammasome. This underscores that while VbP is able to activate the CARD8 inflammasome and cause issues with cytotoxicity, lower concentrations of VbP do not have the same issues and are specific to killing HIV-1 infected cells and enhancing NNRTI-mediated pyroptosis. We also show that the killing and enhancement by VbP are dependent on CASP1 (**Figure 3G**).

### VbP sensitization of the CARD8 inflammasome can overcome NNRTI resistance

As HIV-1 has a high mutation rate, the circulating pool of HIV-1 strains shows distinct genetic variation across clades^35^. The initial work demonstrating protease cleavage of CARD8 demonstrated that proteases from all clades can induce the CARD8 inflammasome, albeit at varying levels of efficiency^8^. This poses a significant barrier for implementation in the clinic as not every patient will have a viral reservoir that is highly sensitive to NNRTI-induced killing. Therefore, we aimed to evaluate this combination strategy against clinical HIV-1 isolates from clades A, B, C, and D. Briefly, CD4^+^ T cells were infected until 10-20% infection was reached, where cells were then treated with EFV or combination along with entry inhibitor T-20 and integrase inhibitor Raltegravir to prevent further rounds of replication. As can be seen in **Figure 4A** each strain demonstrated enhancement upon addition of 0.5 μM of VbP.

**Figure 4:**
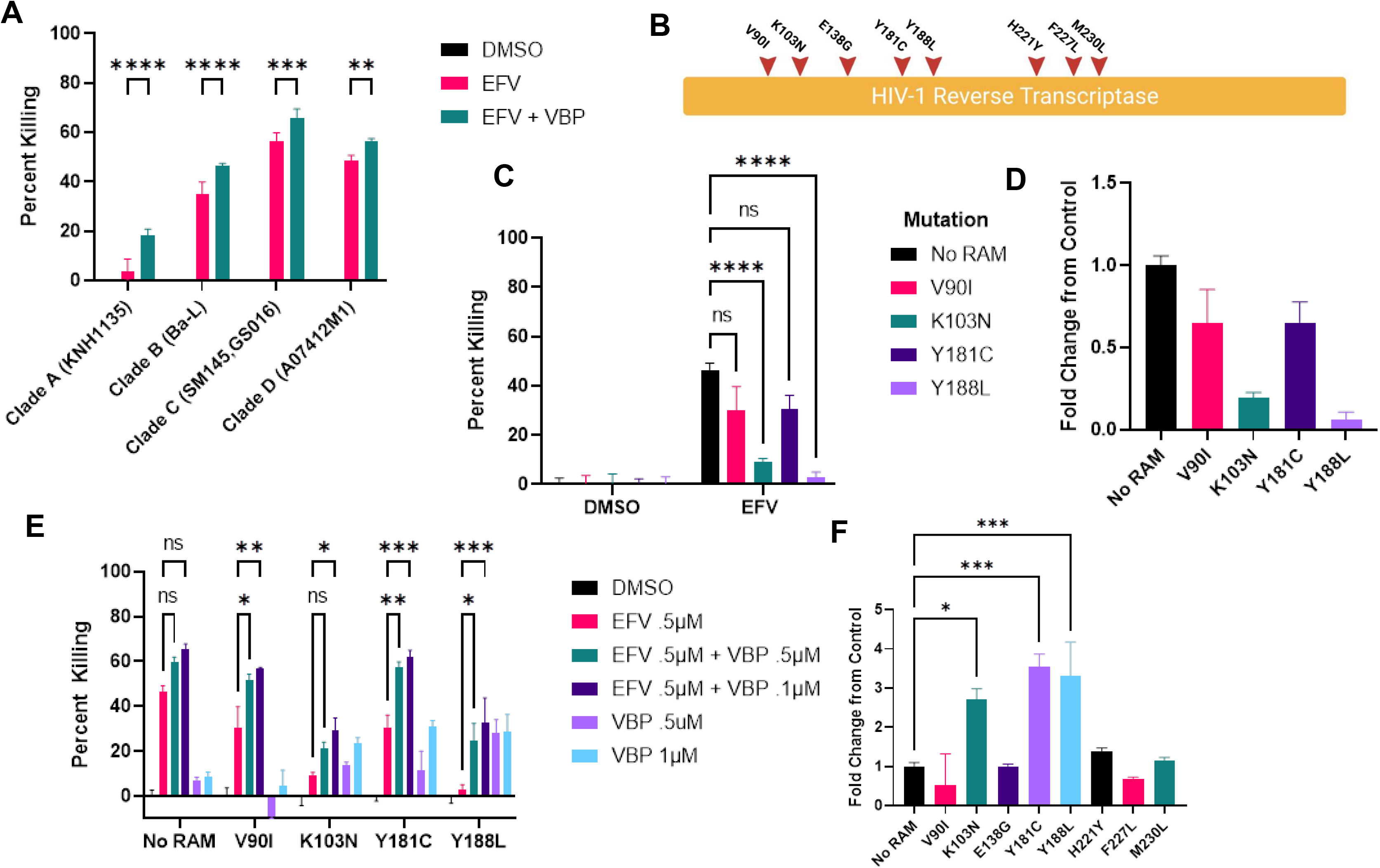
VBP sensitization of the CARD8 inflammasome can overcome NNRTI resistance. **A)** Killing and enhancement of clinical isolates from Clades A, B, C, and D. One Donor of primary CD4^+^ T Cells were treated with DMSO, EFV (.5μM), or EFV with VBP (.5μM) in the presence of media containing T-20 (1μM), Raltegravir (1μM). Each strain showed significant enhancement upon addition of VBP (Two-way ANOVA with Tukey’s multiple comparison test). **B)** Graphical depiction of the location of NNRTI RAMS. **C)** NNRTI RAMS show diminished capacity for killing HIV infected primary CD4^+^ T Cells from one donor upon treatment with .5μM of EFV with respective fold change from no RAM control shown in panel **D** (Two-way ANOVA with Tukey’s multiple comparison test). **E)** Upon VBP treatment, these same mutants show significantly enhanced killing with increasing VBP concentrations (One-way ANOVA with Tukey’s multiple comparison test). Additionally, some RAM containing viruses have increased levels of killing efficiency as depicted by the fold change of killing by 1μM VBP treatment from no RAM virus in panel **F** (Two-way ANOVA with Dunnett’s multiple comparison test). * = p<.05, ** = p<.01, *** = p<.001, **** = p<.0001.

A major concern for implementation of NNRTIs in a “shock and kill” approach is the presence of NNRTI resistance associated mutations (RAMs). As NNRTI RAMs show significant shifts in the EC_50_ values for blocking reverse transcriptase activity, we wanted to first understand if these RAMs would also show resistance to NNRTI-induced CARD8 inflammasome activation^36–39^. RAMs were introduced into our HIV-1 reporter virus (pNL4-3-pol) via site-directed mutagenesis. RAMs were chosen for key regions on HIV-1 reverse transcriptase which can be seen in **Figure 4B**. A set of RAMs were selected for initial testing (V90I, K103N, Y181C, and Y188L) and reductions in killing efficiency compared to the control virus can be seen for all four mutants tested (**Figure 4C**). Although V90I and Y181C did not show statistically significant reductions, p-values in comparison to control were .051 and .053 respectively. Alternatively, K103N and Y188L showed stark, statistically significant reductions in killing efficiency and the fold changes from control can be seen in **Figure 4D**. As previously documented for blocking reverse transcriptase activity, NNRTI RAMs differ in the level of resistance that they confer. Strong NNRTI RAMs may confer complete resistance to NNRTI-mediated killing whereas others may simply show reduced efficacy. This may help in the classification of viral strains that may respond to NNRTI treatment alone versus those that require VbP enhancement for their function. We then tested these NNRTI RAMs for their killing efficiency upon addition of VbP.

While all RAMs tested showed increased rates of killing upon combination treatment, two distinct stories again arise when comparing RAMs with reduced efficacy in comparison to those with complete resistance (**Figure 4E**). The NNRTI RAMs that showed reduced efficacy, V90I and Y181C, demonstrated enhancement with VbP that surpassed the killing of the non-mutant control with EFV alone indicating a significant rescue of killing efficacy. The mutants that conferred near-complete resistance, K103N and Y188L, demonstrated significant increases in killing efficiency with combination treatment compared to EFV alone. However, this enhancement was not significantly different from VbP alone treated samples. As can be seen in **Figure 4F** some viruses are more susceptible to killing by VbP alone such as these mutants that confer complete resistance to NNRTIs.

## Discussion

Since NNRTIs offer a promising strategy for eradication of HIV-1 latent reservoirs, improving their *in vivo* cell killing potency is essential to the treatment efficacy. This study proves that sensitization of the CARD8 inflammasome through DPP9 inhibition can reduce the threshold and provide more effective clearance of HIV-1 infected cells for clinically relevant scenarios. Additionally, we show that DPP9 inhibition through chemical means such as with VbP can induce targeted cell killing on their own which varies across viral strains with NNRTI RAMs. We therefore posit that there may be varying levels of intrinsic Gag-Pol dimerization of these strains which can be enhanced by VbP, which are not due to NNRTIs due to the lack of dimerization induced from NNRTI binding. This suggests that viral strains that confer near-complete resistance to NNRTIs may still be sensitive to targeted killing through the sensitization of the CARD8 inflammasome. Taken together, these data prove that although NNRTI resistance may prove to be a significant barrier in implementation of NNRTIs for a shock and kill approach, they are not insurmountable when the CARD8 inflammasome is sensitized through DPP9 inhibition

The major hurdle for implementation of VbP as a combination therapy is the cytotoxicity at high concentrations. This is likely in large part due to lack of specificity for inhibiting the DPP9 and CARD8 interaction. With the advent of our previous work on CARD8 sensing of HIV-1 protease activity and this work on enhancement of sensing through DPP9 inhibition there is sufficient rationale for the development of new DPP9 inhibitors. Further work will be done to screen for chemical inhibitors that can specifically prevent CARD8C capture rather than binding to the enzyme active site on DPP9, whereas screens for CARD8 inflammasome inducers have great potential for identifying less specific and more toxic compounds. Screening for inhibition of CARD8C capture should hopefully increase specificity for DPP9 inhibition of CARD8 and eliminate the possibility of induction of non-HIV-1-specific CARD8 N-terminal degradation. Through this there should be increased potency and specificity for HIV-1-infected cells leading to more robust enhancement of NNRTI-based strategies and less cytotoxicity due to spurious CARD8 inflammasome activation. While the exact mechanism of VbP induction of the CARD8 inflammasome and enhancement of HIV-1 sensing is still not fully understood, this provides a promising new angle for combination therapy. Additionally, this study provides a proof of concept for in vitro DPP9 inhibition, but further work will be done to assess the efficacy of VbP and EFV combination treatment in vivo.

## Materials and Methods

### Plasmids

Plasmids for replication incompetent viruses used in this study can be found in **Figure S1A**. To generate plasmids for these viruses, mutations were introduced into the pNL4-3-GFP vector (AIDS Reagent Program #111100), which contains an enhanced green fluorescent protein (EGFP) inserted into *env*. L40C-CRISPR.EFS.PAC (Addgene #89393) and SGL40C-H1.EFS.RFP657 (Addgene #69148) vectors were used for sgRNA delivery via lentivirus. CRISPR/Cas9 guide RNAs were selected using the CCTop selection tool (43). pLKO.1puro (Addgene #8453) was used for gene knowckdown via lentivirus vectors. Site-directed mutagenesis to obtain NNRTI RAM’s was done using PCR primers on the NL4-3-Pol plasmid and were confirmed by sequencing; primers can be found in **Table S1**.

### Cell Culture

HEK293T (CRL-3216) and THP-1 cells (TIB-202) were ordered from ATCC and cultured in DMEM or RPMI 1640 medium with 10% heat-inactivated fetal bovine serum (FBS), 1 U/ml penicillin, and 100 mg/ml streptomycin (Gibco). CD4^+^ T cells from blood were isolated from healthy donor peripheral blood mononuclear cells (PBMCs) using the BioLegend human CD4^+^ T cell isolation kit (BioLegend #480010). Purified CD4^+^ T cells were co-stimulated with plate-bound CD3 antibody (Biolegend #300333) with media containing soluble CD28 (Biolegend #302943) antibody and 20 ng/ml IL-2 (Biolegend #589106) for 3 days. Human serum containing media comprised 50% human serum obtained from Gemini Bio (#100-110) with 10% FBS, 1U/mL penicillin, 100mg/mL streptomycin, and 40% RPMI 1640 medium. For MTS assays the CellTiter 96® AQueous One Solution Cell Proliferation Assay (MTS) from Promega was used (Promega #G3580). Uninfected CD4^+^ T cells or THP-1 cells were treated with EFV, RPV, VbP, or DMSO for two days prior to addition of MTS reagent, MTS reading was done following manufacturer’s protocol.

### Preparation of HIV-1 and lentivirus stocks

Reporter viruses were packaged by co-transfecting HEK293T cells with viral vectors, packaging vector pC-Help (44), and pVSV-G (Addgene #8454). To expand clinical HIV-1 isolates, CD8-depleted PHA-stimulated PBMCs were infected with the international HIV-1 isolates (AIDS reagent program #11412). Culture supernatant was collected after 6-9 days and filtered prior to use. Lentiviruses for knockdown or knockout were also packaged in HEK293T cells by co-transfecting pVSV-G, psPAX2 (Addgene #12260), and sgRNA or shRNA using Lipofectamine 2000 (Thermo Fisher). Lenti-X Concentrator (TaKaRa #631232) was used to concentrate supernatant containing virus.

### Generation of THP-1 cells with gene knockout or knockdown

The sgRNA and shRNA sequences can be found in **Table S2** and were verified by sequencing. THP-1 cells were transduced with sgRNA or shRNA lentiviruses via spin inoculation for 2 hours at 1200g at 25°C. Cells were then selected with puromycin (1 μg/ml) for 5-7 days prior to infection with HIV-1 reporter virus NL4-3-ΔVif-Vpr. Immunoblotting was performed to confirm knockout or knockdown efficiency. The controls for knockout cells were transduced with a Cas9-expressing lentiviral vector without sgRNA.

### HIV-1 infection and cell killing

HIV-1 p24 ELISA was used to verify viral stock concentration (XpressBio #XB-1000). HIV-1 reporter virus infection was performed at a multiplicity of infection (MOI) of 10 and 0.1 for clinical isolates. Infection was again performed by spin inoculation (1,200g) for 2 hours at 25 °C. Antiretrovirals (ARVs) were obtained from the NIH AIDS Research and Reference Reagent Program: rilpivirine (RPV), efavirenz (EFV), etravirine (ETR), nevirapine (NVP), T-20, and raltegravir (RAL). Doravirine (DOR), Val-boroPro (VbP), along with additional EFV and RPV, were obtained from Selleck chem(#S6492, #S8455, #S4685, and # S7303). NNRTIs alone or in combination with VbP were added to HIV-11-infected cells 3-4 days post infection. For dose response curves NNRTI’s were serially diluted 3-fold prior to addition to infected cells. For GFP-reporter viruses, infection was determined by flow cytometry. For clinical isolates, intracellular HIV-1-p24 staining was performed using the Cytofix/CytopermTM kit (BD #554714) using anti-HIV-1 p24-PE antibody (#6604667, 1:1000 dilution) purchased from Beckman Coulter. The FLICA660 Caspase1 staining reagents were purchased from ImmunoChemistry Technologies (#9122). Percent infection (GFP+ or p24+) was determined by flow cytometry (BD LSRFortessa, BD FACSCanto, or BD accuri c6 plus). Flow cytometry data were analyzed by Flowjo software.

Percent killing, log fold change in EC50, and fold change of enhancement were calculated as follows:

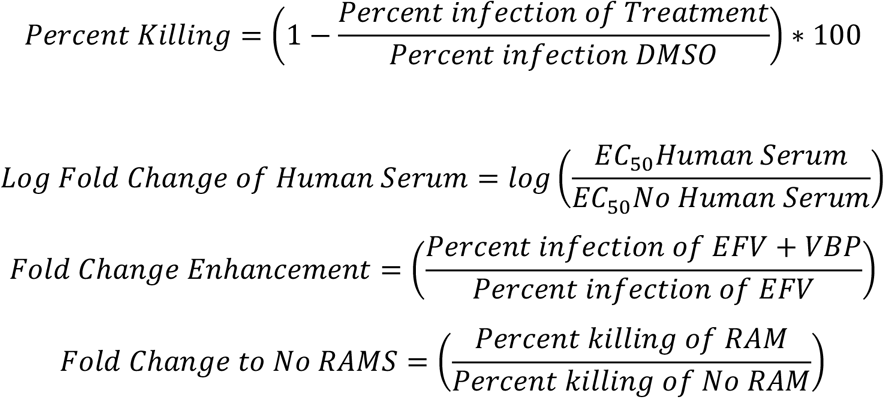

### Statistical Analysis

Statistical analyses were performed using Prism 8 (GraphPad). The methods for statistical analysis were included in the figure legends. Error bars show mean values with SEM. The web application for SynergyFinder2.0 was used to calculate synergy and the output graphs were used. Heatmaps were generated using Python with the Seaborn and Matplotlib packages.

## Acknowledgements

The following reagents were obtained through the AIDS Research and Reference Reagent Program, Division of AIDS, NIAID, NIH: rilpivirine, efavirenz, lopinavir, etravirine, nevirapine, maraviroc, T-20, tenofovir, raltegravir, pNL4-3-GFP, international HIV-1 isolates, and HIV-1 p24 antibodies. Funding: This work was supported by NIH grants R01AI162203 (to L. Shan). Author Contributions: L.S. and K.C. designed the study, analyzed the data, and wrote the manuscript. K.C. and Q.W. performed the experiments. Competing interests: the authors declare no competing financial interests.

