## Supplement figures and tables for "CARD8 inflammasome sensitization through DPP9 inhibition enhances NNRTI-triggered killing of HIV-1-infected cells"

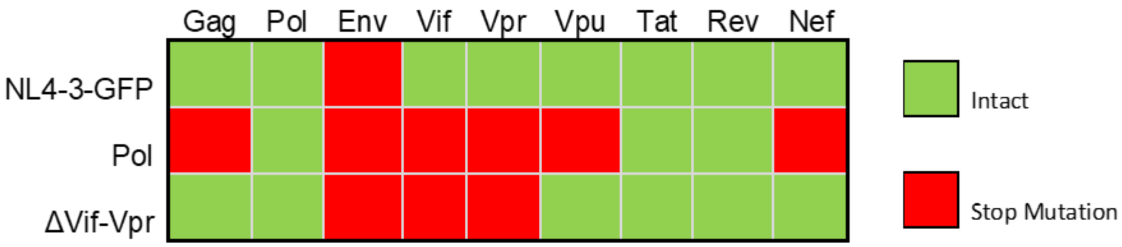


**Figure S1: NL4-3 Reporter Viruses**
Three main EGFP reporter viruses were used for this study: NL4-3-GFP, NL4-3-Pol, or NL4-3-Δvif-vpr. NL4-3-pol contained truncations in all genes except pol, tat, and rev and was used for all $CD4^+$ T Cell experiments. NL4-3-GFP was used for initial THP-1 dose response, but all other THP-1 experiments used the NL4-3-Δvif-vpr due to lower toxicity and better cell survival using this virus.

| Primer Name | Sequence (5'-3') |
|---|---|
| Fragment_1_F | gacatagcaggaactactagtacccttcaggaacaaatagg |
| Fragment_2_R | aatacactccatgtaccggttcttttagaatctccctg |
| V90I_1_R | taattgaatttcccagaaatcttgagttctctt |
| V90I_2_F | tgggaaAttcaattaggaataccacatcctgca |
| K103N_1_R | gatttgttctgttttaaccctgcaggatgtgg |
| K103N_2_F | aaaacagaacaaatcagtaacagtactggatgtgg |
| E138G_1_R | ccctggtgtcccattgtttatactaggtatggt |
| E138G_2_F | cctagtataaacaatgggacaccagggattaga |
| Y181C_1_R | catgtattgacagatgactatgtctggattttgttt |
| Y181C_2_F | atagtcatctgtcaatacatggatgatttgtatgta |
| Y188L_1_R | agatcctacaagcaaatcatccatgtattgatagat |
| Y188L_2_F | gatgatttgcttgtaggatctgacttagaaataggg |
| H221Y_1_R | ttctttctgatatttttttgtctggtgtggtaaa |
| H221Y_2_F | acaccagacaaaaaaatatcagaaagaacctcca |
| F227L_1_R | ataacccatccaaaggagtggaggttctttctg |
| F227L_2_F | cctccactcctttggatgggttatgaactccat |
| M230L_1_R | ataacccagccaaaggaatggaggttctttctg |
| M230L_2_F | cctccattcctttggctgggttatgaactccat |

**Table S1: NNRTI RAM primers**
NNRTI Resistance associated mutation primers for site-directed mutagenesis are listed. All mutants used the same F1 and R2 primers listed.

| sgRNA/shRNA | Sequence (5'-3') |
|---|---|
| CARD8 sgRNA | TGAGGCCTTGCTGAGCATGG |
| CASP1 sgRNA | TTTATCCGTTCCATGGGTGA |
| NLRP1 sgRNA1 | GCTCAGCCAGAGAAGACGAG |
| NLRP1 sgRNA2 | TATGTGATGCAGCTCCACCC |
| NLRP1 sgRNA3 | GATAGCCCGAGTGACATCGG |
| NLRP1 sgRNA4 | AGCCCGAGTGACATCGGTGG |
| NLRP1 sgRNA5 | GGAGCAGTACGAGAGGGTGC |
| NLRP1 sgRNA6 | ACACGAGGCCCAGCCAGATG |
| shScramble | CCTAAGGTTAAGTCGCCCTCG |
| DPP8 shRNA1 | GCTGGTGAATAACTCCTTCAA |
| DPP8 shRNA2 | GCTGCACTTTCTACAGGAATA |

**Table S2: sgRNA and shRNA Sequences**

Sequences used for knockdown and knockout of key inflammasome genes are listed above.

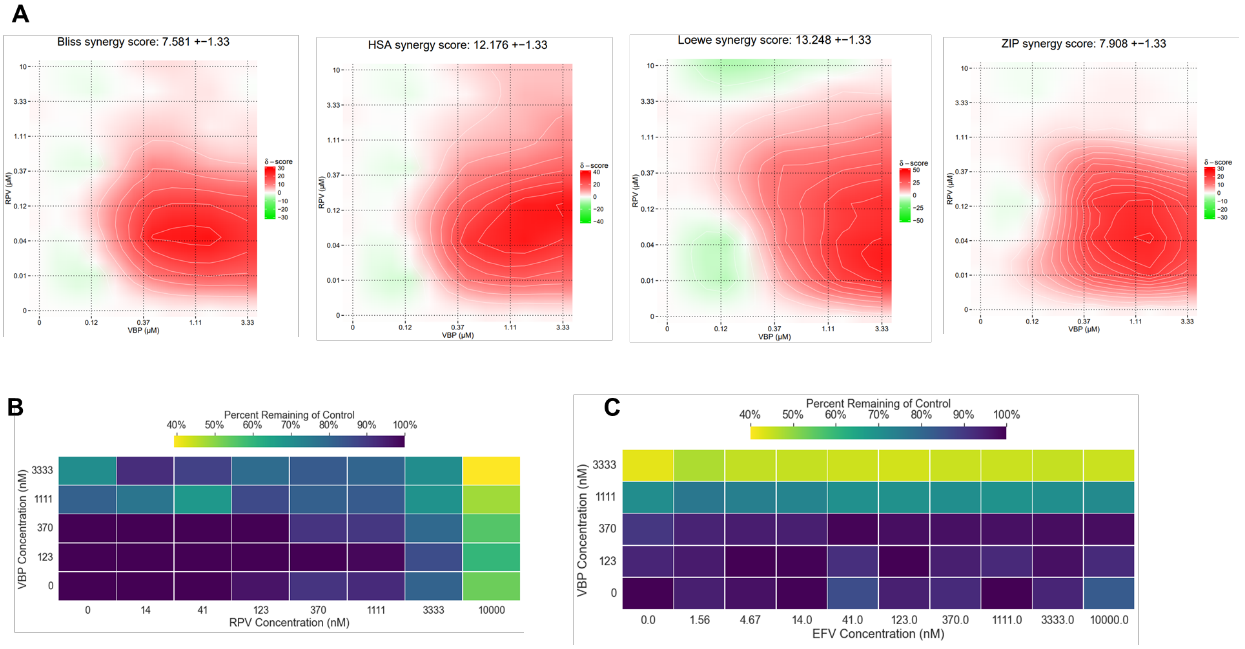


**Figure S2: DPP9 inhibition sensitizes the CARD8 Inflammasome to NNRTI-mediated pyroptosis**

**A)** VBP and RPV combination treatment denotes a synergistic relationship as evidenced by four independent synergy calculation methods from SynergyFinder2.0 (Bliss, HSA, Loewe, and ZIP). The highest levels of synergy are found at sub-micromolar RPV concentrations and concentrations of VBP > 123nM. **B)** VBP is not toxic below 3.33µM in CD4$^+$ T Cells as denoted by the heatmap of MTS assay results of three separate donors of primary CD4$^+$ T Cells treated for two days with RPV and/or VBP, however RPV at high concentrations presents significant cytotoxicity. **C)** VBP is not toxic below 1.1 µM in THP-1 cells as denoted by the heatmap of MTS assay results in THP-1 cells treated for two days with EFV and/or VBP. Increased cytotoxicity in THP-1 cells may be due to higher levels of inflammasome components in these cells.

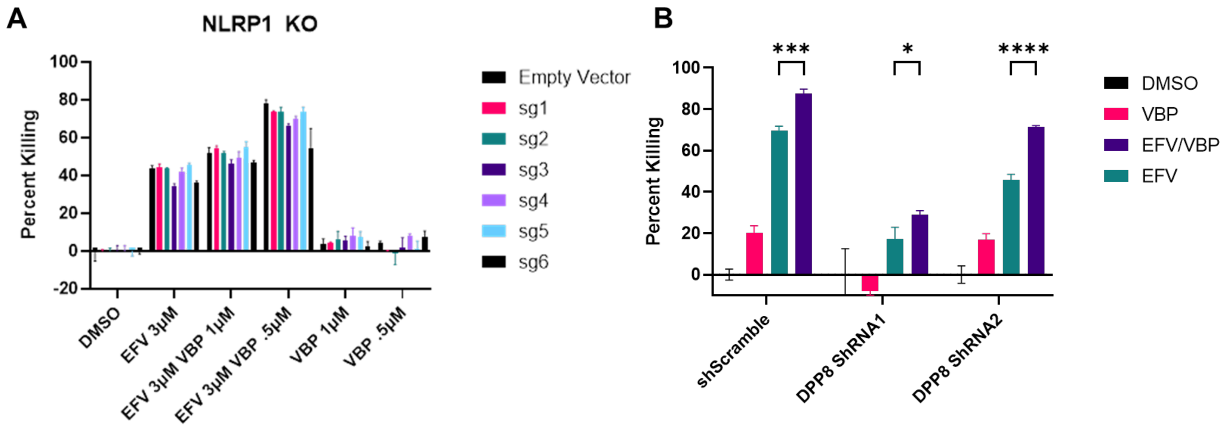


**Figure S3: Characterization of VBP enhancement of NNRTI induced cell killing**

**A)** Upon bulk knockout of NLRP1 in THP-1 cells VBP is still able to enhance killing of HIV-1 infected cells. Six separate guide RNA's were tested for knockout, sgRNAs 1-4 target the FIIND domain, sgRNA5 targets the pyrin domain, and sgRNA6 targets the CARD domain of NLRP1. Regardless of the guide RNA used enhancement is still present indicating lack of involvement of NLRP1. **B)** DPP8 is also not essential for VBP enhancement of NNRTI-mediated killing of HIV-1 infected cells. DPP8 knockdown was done using two separate shRNAs targeting DPP8 in comparison to a scramble shRNA control. VBP enhancement is still significant upon treatment with EFV and VBP together (* = p<.05, *** = p<.001, ****=p<.0001 by two-way ANOVA with Tukey's multiple comparison test).
